# A Practical Approach to Estimating the Effect of Synchrotron Radiation on Sperm Motility and Viability

**DOI:** 10.1101/2023.06.08.544169

**Authors:** Mette Bjerg Lindhøj, Anne Bonnin, Kristian Almstrup, Tim B. Dyrby, Rajmund Mokso, Jon Sporring

## Abstract

Studying the dynamics of sperm tail beating in 3D is important for understanding decreasing fertility trends related to motility. Synchrotron X-ray tomography (SXRT) shows promising results for imaging live biological samples in 3D and time, making it a candidate for imaging sperm tail beating patterns. However, the dose of ionizing radiation (IR) that the cells would receive during image acquisition is of concern as this is likely to affect essential cellular functions. The effect of IR on motility at the dose rates encountered in a synchrotron has never been assessed. Here, results are presented of analyzing the movement and viability of sperm cells exposed at varying durations at the TOMCAT beamline (Swiss Light Source (SLS)) which has the field of view and spatial resolution needed to reconstruct sperm cells coupled with sub-second exposure times. The results indicate that motility is affected long before viability and that sperm cell dynamics are affected very quickly by the beam.

## 1 Introduction

According to Abbe’s diffraction limit, microscopes can not resolve objects closer than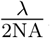, where *λ* is the wavelength of the light source, and NA is the numerical aperture. This naturally allows electron-and X-ray microscopes to have a higher image resolution than optical microscopes, as these light sources have a much lower wavelength. A shallow depth of field further limits standard optical microscopes whereas, electron and X-ray microscopes have the added advantage of allowing 3D imaging. Recent advances in synchrotron X-ray technology and detector systems have opened the possibility for synchrotron X-ray tomography (SXRT) to be used for dynamic 3D imaging of live biological samples (1), which is unique to this modality. However, successfully imaging live specimens is challenging as it requires simultaneously optimizing image quality on temporal and spatial image resolution at a dose where the sample survives.

Dynamic single-cell studies of sperm cells in 3D are relevant for understanding parameters important for motility. Recent studies have shown that the 3D dynamics reveal motion patterns that cannot be directly observed under a standard optical microscope where only the 2D motion is visible (2). Changes in sperm motility caused by environmental chemicals acting on the primary calcium channel of sperm have been proposed as a mechanism involved in recent decreasing fertility trends (3, 4), and synchrotron imaging is a candidate tool to further study the effect of environmental chemicals on single-sperm motility. Within the last decade, image resolution of SXRT imaging has been pushed to the micro-and nanometer scale with sub-second exposure times (5–14). Hence, it could be feasible to perform SXRT of a sperm cell to analyze the tail beat. However, a fundamental question is whether the cells can survive in the beam and whether the tail beating could be unaffected long enough for the data to be relevant.

Radiation can cause severe damage to living organisms, including oxidation, DNA damage, cell death, inactivation of muscle function and incapacitation and death in various insects and small animals (15–23). A previous study coupled dose to image resolution in unlabelled biological samples under cryogenic conditions (24). The study estimated that the dose for unlabelled protein in amorphous ice at a 100 nm image resolution would be close to the dose which is associated with immediate inactivation of muscle function (25) and above the dose where the majority of radiation-resistant bacteria die (18). Typically, SXRT of cells is performed under cryogenic conditions in a unique energy band called the water window, where the contrast of unlabelled cells is optimal, and the damaging effect of reactive oxygen species are minimized. However, labelling could be an option to increase the contrast and allow imaging at higher energies. At higher energies, the attenuation coefficients of soft tissue are lower, decreasing the dose absorbed by the sperm. Furthermore, the impact of radiation damage could be different in sperm, bacteria and complex organisms such as insects and mammals. Radiation effects on male fertility have been studied before (26), but only considering the detrimental effects to spermatogenesis during medical radiotherapy.

The damage caused by ionizing radiation is not only a question of the total dose delivered. The damage takes time to occur, and different types of damage occur at different time scales and do not necessarily end when the exposure stops. Third generation light sources, which produce brilliant X-rays, allow fast imaging by having a very high flux, delivering high doses at a fast rate. Hence, sperm imaged in such a synchrotron would receive a high radiation dose, but whether they would be motile long enough to capture the natural tail beating pattern remains an open question. Additionally, sperm is an exciting subject for probing the effects of radiation damage on cells as they are among the most dynamic cells and readily available, allowing us to measure changes to both viability and motility.

Dynamic single-cell studies pose many practical problems in terms of sample preparation. Catching and containing a single cell within the field of view can be very difficult as perishable live samples must be prepared on-site with available or portable equipment, and these differ between instruments. The experimental method described in this article provides a practical approach to assess the damage caused by a specific beam on a population of sperm and estimate the relative dose on a single-cell sample. The advantage is that the population study is much easier to perform and can therefore be used to probe the feasibility of a single-cell experiment.

## 2 Methods and Material

To better understand how synchrotron radiation affects sperm, a test was at the TOMCAT beamline (Swiss Light Source (SLS)). A population of sperm were exposed to the X-ray beam while simultaneously monitoring their movement with an optical microscope camera. Subsequently, a NucleoCounter^®^ SP-100™cell counter was used to assess the viability of the sperm. In the following, the experimental setup is explained. Then the analysis of the video data to estimate the effect on sperm motility is presented. Finally, as the experiment should indicate the feasibility of a possible live single-cell experiment, and the dose is not directly comparable, an estimation of the dose relationship between the performed population experiment and a future single-cell experiment is presented. The estimate is based on using the Nanoscope, a full-field transmission X-ray microscope (TXM), where the X-ray beam is focused with optics to obtain the required resolution and field of view for single-cell imaging.

### 2.1 Mounting a video camera above an exposed sample can be used to assess sperm motility

A population experiment was performed to assess the effect of the X-ray beam on the sperm. We placed 1 mL of Duroc boar sperm in a Ø 3.5 cm plastic Petri dish. The Petri dish was positioned with laser on a mirror on rotation stage in front of the X-ray beam. Above the dish, we mounted an optical microscope camera, see Figure 1. The mirror increased the light needed to see the cells. The dish was placed so that the X-ray beam would go approximately through the centre of the dish.

**Fig. 1.**
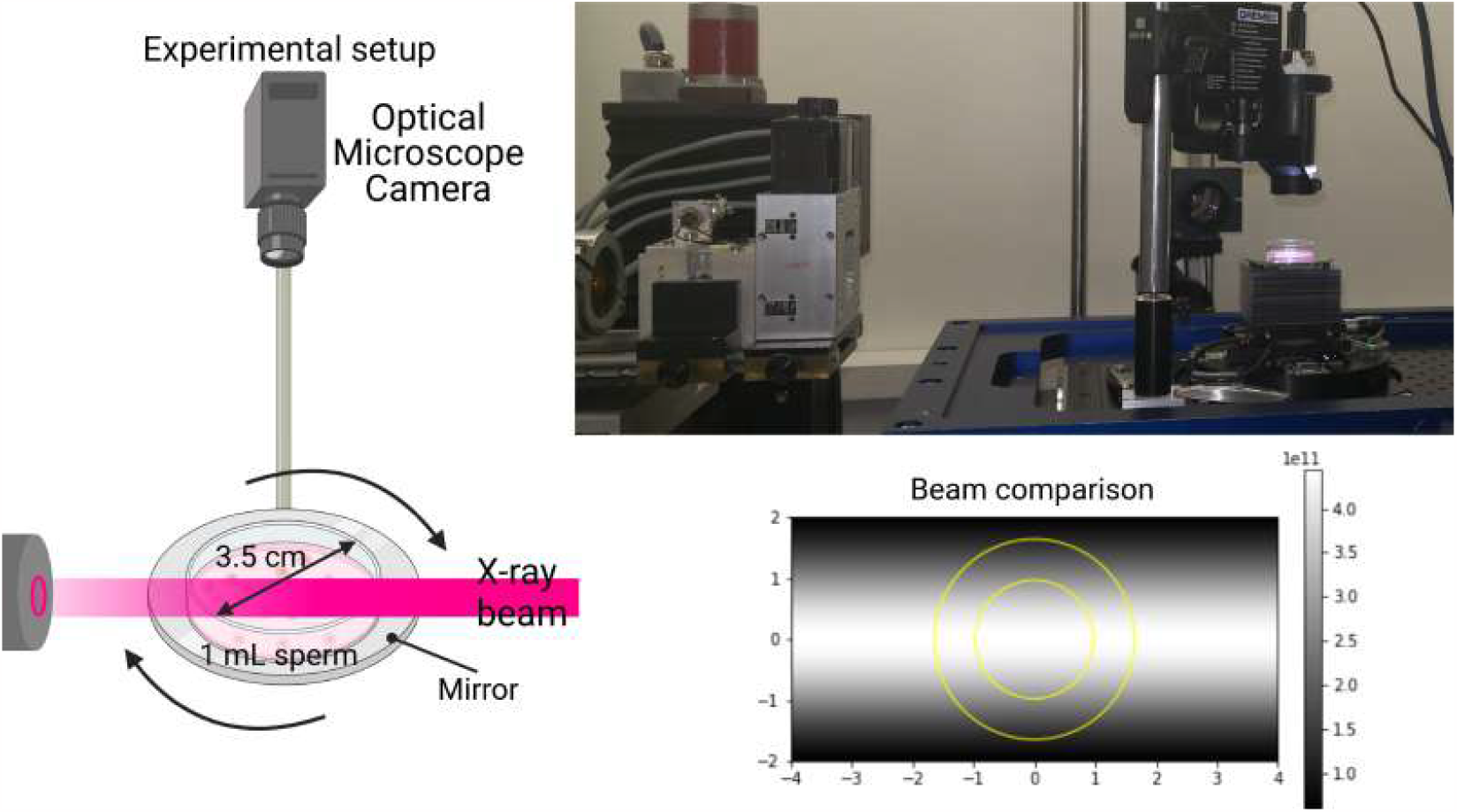
Experimental Setup. On the left is a schematic drawing of the experimental setup with the camera mounted above an Ø3.5 cm petri dish with 1 mL sperm in it. On the top right is a picture of the actual setup. The bottom right is a plot of the X-ray beam model used in the dose calculation. The background image shows the X-ray beam intensity profile as a colour gradient with white being *N*_*photons*_ and black being no photons. The yellow doughnut on top is the part of the X-ray beam used by the Nanoscope. The figure was made using BioRender.com.

We recorded a video with the optical microscope camera, which was left to record before, during, and after X-ray exposure. We performed five experiments at varying X-ray exposure times. The sperm in the five tests were exposed for 1,5,10,15, and 45 minutes. The Petri dish was turned on the rotation stage to expose all cells as evenly as possible during exposure. Immediately after X-ray exposure, we measured the number of dead cells in a NucleoCounter using a membrane-impermeable DNA intercalator, propidium iodide.

The full X-ray beam was used to maximize the number of cells exposed to the beam during the experiment. The Nanoscope X-ray beam, which can be used to image single sperm cells, is focused by optics to obtain the required resolution for single-cell imaging and is therefore different from the beam used for the performed motility test, see Figure 1. The difference affects the dose, which is analyzed below. The full X-ray beam used during the experiment was 8 mm wide and 4 mm high.

A fresh sperm sample was drawn from the same batch during the whole experiment for each test. The tube was gently turned a couple of times before the sample was taken to distribute the cells well. During X-ray exposure, the Petri dish was turned using the rotation stage. Before counting the cells in the cell counter, they were gently mixed with the pipette to disperse them in the liquid uniformly. All measurements were done twice.

The cell counter measured the cell density before and after the experiment. As a control, sperm not exposed to the X-ray beam was used. The control eliminated the possibility that cells were dying due to other factors.

### 2.2 Entropy and correlation measures between frames can be used to assess the motility in the recorded videos

The videos recorded during the experiment were analyzed to assess the damage to the motility of the sperm cells. We used two measures; the correlation of the intensity values between frames and the entropy of the pixels in the time direction.

The videos were loaded into python using the *av* library. The videos were in HSV format, and we only considered the last channel. We analyzed 100 frames prior to X-ray exposure and 100 frames after X-ray exposure. The number 100 was chosen because 100 frames were available before and after X-ray exposure in each video. The dimensions of the resulting data blocks were 100 *×*1280 *×*720 pixels. The resulting blocks were a 3D dataset with two spatial axes and a time axis. The spatial dimensions were flattened to get a 2D matrix with the pixel intensity varying with time through each column. We determined the entropy value within each column using the scikit-image (27) local entropy function with our 2D matrix and a neighbourhood corresponding to a single column. The median of the entropy over all pixels was chosen as the final entropy value. Numpy was used to calculate the Pearson product-moment correlation coefficients between frames.

### 2.3 The dose relationship between a population experiment and a single-cell setup can be estimated

The dose relationship between the performed experiment and a future singlecell experiment must be estimated to assess the feasibility. For the estimate, we will use the equation for skin entry dose of (28)

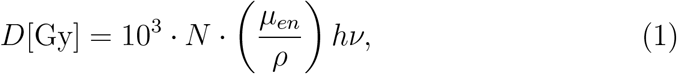

where *D* is the skin entry dose, N is the total number of incident photons per cm^2^, ^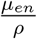^ is the mass energy absorption coefficient given in cm^2^ g^*−* 1^ which can be obtained from the NIST database (29) and *hν* is the energy of one photon in J.

The terms in (1) will be compared starting from the last term to make a comparison between the skin entry dose in the performed population experiment and the skin entry dose in a future single-cell experiment. The population experiment was performed at 10 keV, which is the same energy that would be used for a future single-cell experiment. Hence *hν* would not change. The term 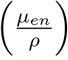 is dependent on the material in the sample and therefore would not change. The only difference between the dose in the performed experiment and a future single-cell experiment is *N*, the total number of incident photons per 1 cm^2^. The full X-ray beam was used during the performed population experiment. The Nanoscope X-ray beam would be used for a single-cell experiment. The X-ray beam model for each X-ray beam is displayed in Figure 1.

The number of photons measured at the center of the full X-ray beam is *N*_*photons*_ = 4.4368 *·*10^11^[photons*/*mm^2^ *·*s] at 10 keV. The number of photons is not constant throughout the X-ray beam cross-section. In the horizontal direction, the X-ray beam is close to constant; the beam follows a Gaussian curve in the vertical direction. Hence, the following approximation will be used for estimating the number of photons during one second per mm^2^, *f* (*x, y*), at a point (*x, y*) in the X-ray beam centered at (0.0), see Figure 1:

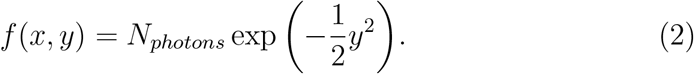

To get the total number of photons *N*_*fullbeam*_ in the X-ray beam, an integration of *f* (*x, y*) over the total area can be performed

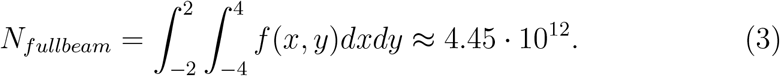

To calculate the number of photons in the Nanoscope beam *N*_*nano*+_, (2) is converted to polar coordinates, the integral over the Nanobeam area is determined

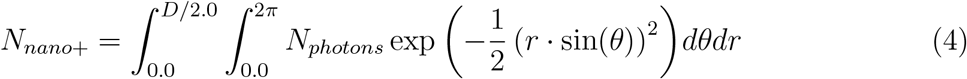

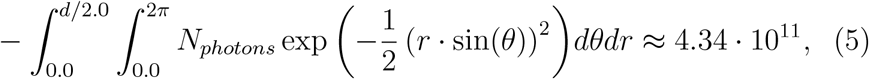

where *D* = 3.294 is the diameter of the outer yellow circle in Figure 1 and *d* = 1.946 is the diameter of the inner circle, both in mm. Furthermore, the Nanoscope uses a condenser which has an efficiency of 20%. Therefore, the total number of photons from the Nanoscope which would reach a single-cell sample is *N*_*nano*_ = 0.2*·* 4.34*·* 10^11^ = 8.68 *·* 10^10^.

By comparing the two estimates, we can estimate that the total number of photons entering the Petri dish in the performed experiment is approximately 50 times higher than the number of photons that would reach a single-cell sample.

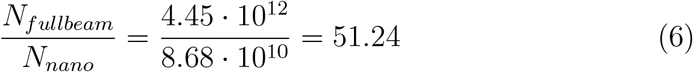

An X-ray beam is attenuated through a volume according to the Lambert-Beer law. The law can estimate the photon count at the centre of the Petri dish.

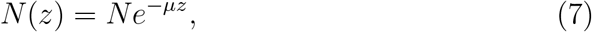

where N is the number of photons, z is the position in the direction of the beam in centimetres, and *µ* is the attenuation coefficient in cm^*−* 1^. The petri dish is very thick, which means that the dose at the X-ray beams exit point of the petri dish is much lower than the entry point. The number of photons reaching the centre of the Petri dish will be calculated to account for the beam attenuation.

The sperm swim in a buffer with similar attenuation properties to water. The sperm will be approximated by soft tissue in terms of x-ray attenuating properties. According to the NIST database, liquid water has a mass attenuation coefficient of 5.329 cm^2^ g^*−* 1^ and soft tissue has a mass attenuation coefficient of 5.379 cm^2^ g^*−* 1^ at 10 keV. They are very similar and exhibit similar x-ray attenuation properties. Therefore, the medium will be approximated as a uniform soft tissue mass. The density of soft tissue according to NIST is 1.060 g cm^*−* 3^, giving *µ* =5.649 cm^*−* 1^. Therefore, the number of photons reaching the centre of the dish is much reduced:

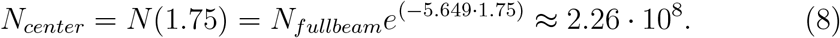

A single-cell sample would only be 75 µm in diameter, and therefore the absorption would be close to zero for such a sample. Comparing the above result with the photon count in the Nanoscope beam, it becomes clear that at the centre of the Petri dish, the number of photons is approximately 380 times smaller than the number of photons in the Nanoscope beam:

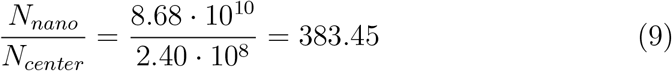

As the sperm were imaged approximately at the centre of the Petri dish, this value is more indicative of the dose relationship. It is important to note that the rotation of the Petri dish affects the radiation dose, and it is not considered here. The calculations presented are meant to serve as a rough estimate.

## 3 Results

### 3.1 Viability of sperm at various exposure times showed a significant effect at 45 minutes exposure

We exposed the sperm in the Petri dish for five different durations, 1, 5, 10, 15 and 45 minutes. After the exposure we measured the sperm viability in a NucleoCounter^®^ SP-100™which has an advised measurement range of 0.5 Mio. mL^*−* 1^ to 7.0 Mio. mL^*−* 1^. At exposure times 1,5,10, and 15 minutes the number of dead sperm did not change significantly and remained below the advised detection limit. The sample subject to 45 minutes of exposure had just above one million dead sperm. As one million is within the advised measurement range and significantly higher than any of the other viability measurements. Hence, we can reliably state that exposure times of 45 minutes damage the sperm membranes. The total number of sperm before exposure was measured with the NucleoCounter to be 20.14 Mio. mL^*−* 1^. Even after 45 minutes of exposure, only 5 % to 6 % of the sperm were dead.

### 3.2 Analysing videos of exposed sperm showed decreasing motility with increasing exposure times

Right after exposure, we visually inspected the sample in an optical microscope to assess the sperm’s motility damage. The sperm in the 1-minute exposure sample were very active and kept moving even after being left on the counter for a prolonged period. After 5 minutes of exposure, approximately half the sperm were moving, and the sperm motility steadily declined when left on the counter, indicating that the radiation, in addition to an acute decrease in motility, also had some long term effect on sperm motility. The sperm motility was substantially reduced above five minutes of exposure, and no moving sperm could be found after forty-five minutes.

To quantify these changes more precisely, we analysed five videos of the sperm during each of the five exposure times. The microscope camera, a Dino-Lite AM5018MZTL that we used to record the videos, had a large field of view but a low image resolution. The sperm are seen as small bright spots in the videos^1^, and single sperm cannot be distinguished. However, from the videos, it is clear to see the bright flickers of the sperm as they move. The flickering can also be seen by visually inspecting slices in the temporal direction through the video clips, where the sperm appear as bright streaks in the time directions, see Figure 2. Before exposure, all the slices look similar, with the streaks shifting in the spatial direction over time. The streaks gradually appear as more straight vertical lines with minimal shifting in the spatial direction, with increasing exposure times. The increasing uniformity of the streaks indicates less active sperm.

**Fig. 2.**
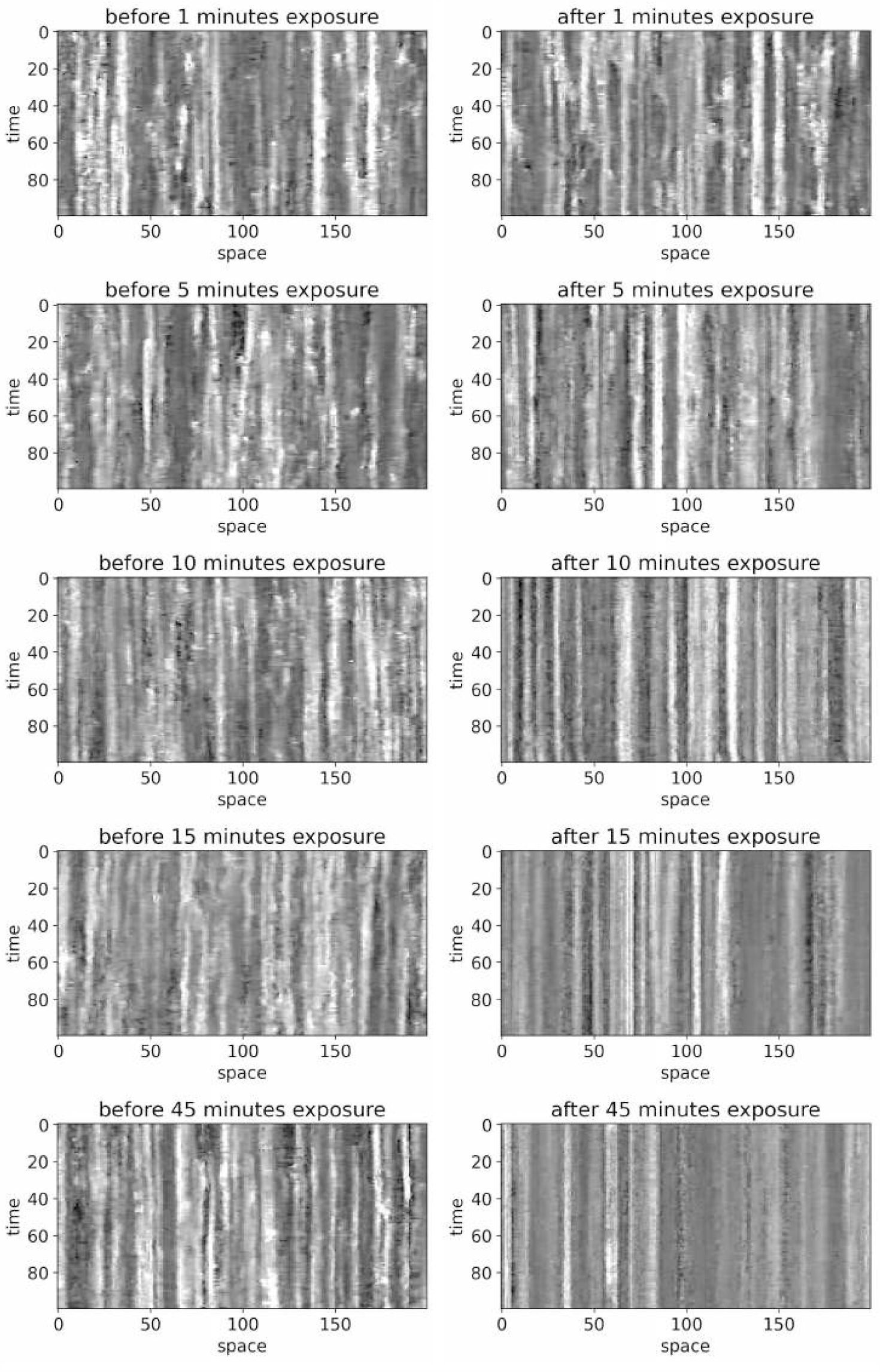
Temporal slices: A table of temporal slices trough each of the video clips before and after exposure (rows) at different exposure times (columns). Looking at the columns of each image, it is clear that the intensity bands become more constant with longer exposure times.

To quantify the sperm motility, we measured the correlation between the frames and the median entropy of all the pixels in the temporal direction, see Figure 3 and Figure 4. The histograms of the correlation values between individual frames in the video clips shift towards higher values after exposure, and the spread of the values decreases, showing that the frames become increasingly correlated after exposure. As exposure time increases, the effect increases, consistent with motility decreasing with increasing exposure time. The entropy measurement shows that the entropy is similar in all video clips before exposure. After exposure, all entropy values are lower than before exposure, indicating that the pixel intensities are less random in the time directly after exposure. We also see that the entropy decreases with longer exposures, meaning less overall flickering of the pixel values for higher exposures. When fitting a linear model to the entropy values after exposure, we obtain an *r− value* of *−* 0.91. The intercept of the linear fit is 0.03, which is close to the median entropy value of all videos before exposure, as expected.

**Fig. 3.**
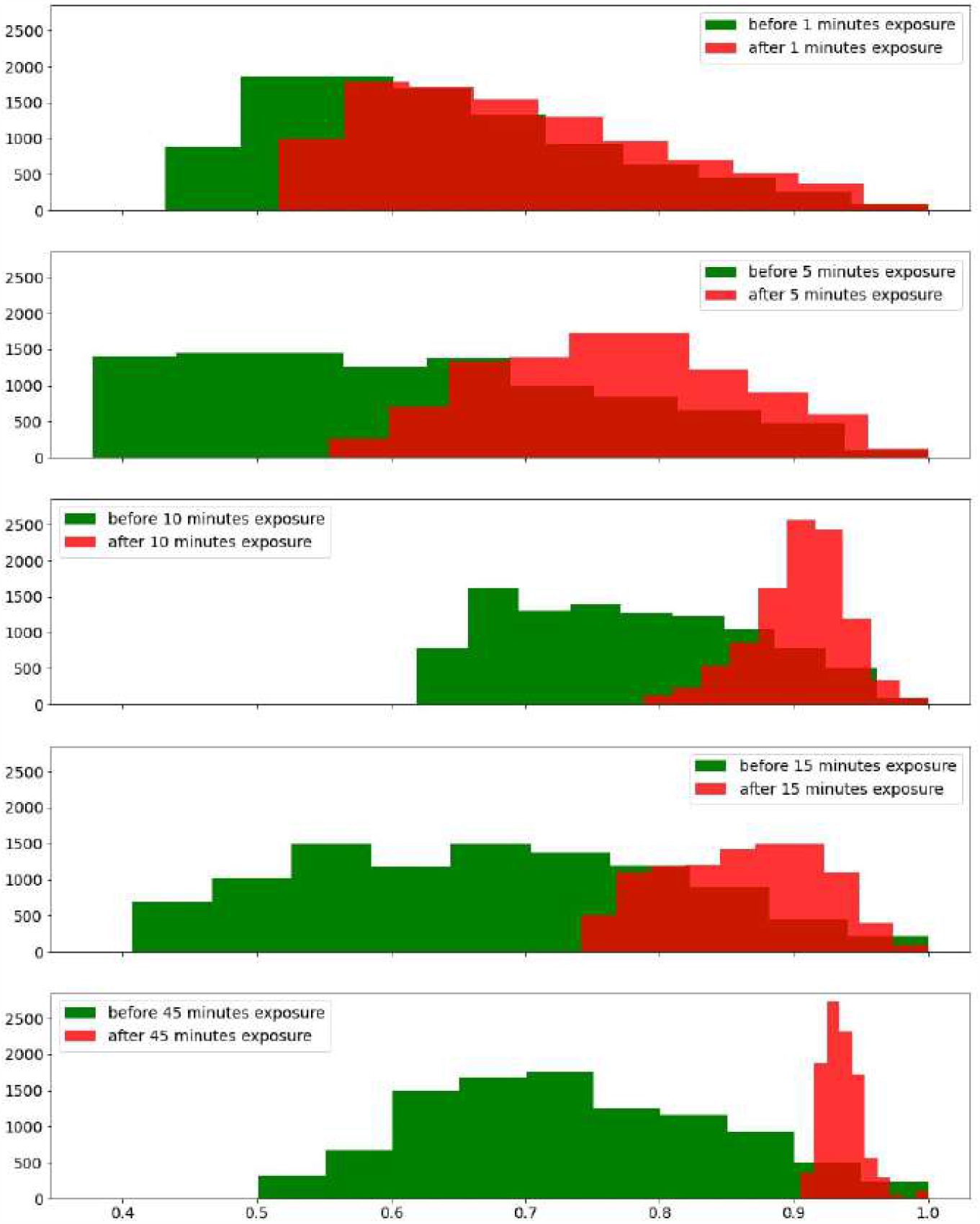
Autocorrelation between frames. Histograms of the correlation values between the individual frame vectors in the selected portion of the movies. We observe that as exposure time increases (down) the spread of the correlation values after exposure (red) decreases and values move to the right (higher).

**Fig. 4.**
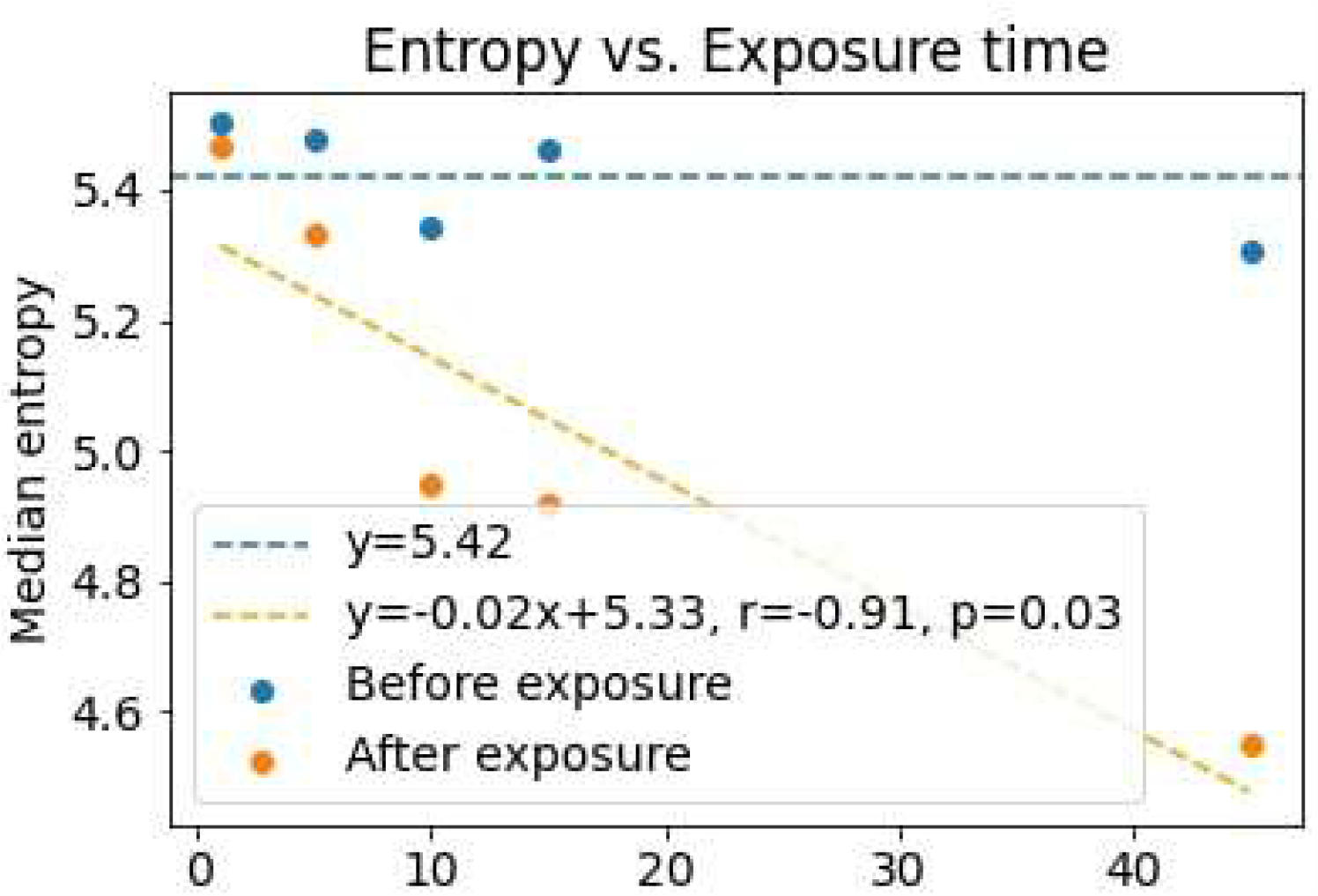
Entropy. The plot shows the median entropy values measured from the five movies. The blue values are from the frames before exposure; the orange values are from the frames after exposure; see Materials and Methods for details. The dashed blue line is a horizontal line corresponding to the median entropy of all videos before exposure. The dashed orange line is a least-squares fitting to the median entropy values after exposure.

## 4 Discussion

Our results suggest that sperm motility is affected much before viability when exposed to the X-ray beam. To the best of our knowledge, the direct effect of radiation damage to the sperm themselves has never been studied. The videos’ correlation and entropy analysis clearly show that sperm motility decreases with increased exposure. At 45 minutes of exposure, the sperm do not move while only 5 % to 6 % of the sperm are classified as dead using the NucleoCounter. The damage caused to sperm by ionizing radiation could be related to several different events. The ionizing radiation could directly affect sperm motility by uncoupling the energy production in mitochondria or causing radiolysis in the surrounding fluid, resulting in damage from free radicals produced in this process. When free radicals accumulate, oxidative stress occurs, which can affect cell signaling (30) and subsequently cause sperm to stop moving, even before any membrane damage has occurred. Furthermore, it seems that cell death measured by membrane permeability is a much slower process and measured in the order of minutes (31).

The radiation dose during the experiment used for assessing the effect on motility was not directly comparable to a future single-cell experiment. During the population experiment, the full X-ray beam was used to expose as many cells as possible and quantitatively assess the damage done by the beam. However, to use the same instrument for SXRT, the focused Nanoscope beam would be used. The dose calculation indicates that a single-cell would receive a higher dose than the cells imaged in the centre of the petri dish during the performed population experiment. The dose calculations do not include sample rotation or the fact that the Petri dish was not fully illuminated as a single-cell sample would be, hence to get a more accurate estimate, a method for including these factors should be found.

The video data is subject to some variability. Known sources include placement of the petri dish and alignment of the microscope, which were both done manually and hence are prone to inaccuracies. Furthermore, the stage is rotated during the exposure, so the area captured by the camera is not identical before and after exposure. Given these variabilities and the small number of different exposure times (5), more measurements are needed to determine an exact relationship between dose and motility.

In conclusion, exposure to ionizing radiation from the full X-ray beam affects sperm motility within a few minutes, while cell viability only seems affected after X-ray extended exposure. The experimental setup described in this article provides a way to assess the radiation damage to sperm function right after exposure before long term effects take over. As sperm are among the most dynamic cells known and easy to come by, the method provides a practical way to assess functional damage caused to sperm by ionizing radiation. As the motility is already affected after short X-ray exposure times, this first attempt shows that it is unlikely the TOMCAT Nanoscope can image the unaffected tail beating of live sperm without changes to the setup. Our method provides a way to estimate the feasibility of future live single-cell SXRT experiments concerning radiation damage.

## Supporting information

publication license from biorender for figure

## 5 Acknowledgement

MBL was supported by the Capital Region Research Foundation (Grant number: A5657) on the MAX4Imagers project (PI: Tim Dyrby).

We acknowledge the Paul Scherrer Institut, Villigen, Switzerland, for the provision of synchrotron radiation beamtime (20201724) at the beamline TOMCAT of the Swiss Light Source.

## Footnotes

1 The videos can be found here: https://data.mendeley.com/datasets/kkszwvx7gk/draft?a=a6a070ec-f148-4fc0-8198-6c0ad3435e95

## Notes

https://data.mendeley.com/datasets/kkszwvx7gk/draft?a=a6a070ec-f148-4fc0-8198-6c0ad3435e95

### Competing Interest Statement

The authors have declared no competing interest.

